# Identification of candidate genes underlying nodulation-specific phenotypes in *Medicago truncotula* through integration of genome-wide association studies and co-expression networks

**DOI:** 10.1101/392779

**Authors:** Jean-Michel Michno, Liana T. Burghardt, Junqi Liu, Joseph R. Jeffers, Peter Tiffin, Robert M. Stupar, Chad L. Myers

**Affiliations:** Bioinformatics and Computational Biology, University of Minnesota; Department of Agronomy and Plant Genetics, University of Minnesota; Department of Plant and Microbial Biology, University of Minnesota; Department of Computer Science and Engineering, University of Minnesota

## Abstract

Genome-wide association studies (GWAS) have proven to be a valuable approach for identifying genetic intervals associated with phenotypic variation in *Medicago truncatula*. These intervals can vary in size, depending on the historical local recombination near each significant interval. Typically, significant intervals span numerous gene models, limiting the ability to resolve high-confidence candidate genes underlying the trait of interest. Additional genomic data, including gene co-expression networks, can be combined with the genetic mapping information to successfully identify candidate genes. Co-expression network analysis provides information about the functional relationships of each gene through its similarity of expression patterns to other well-defined clusters of genes. In this study, we integrated data from GWAS and co-expression networks to pinpoint candidate genes that may be associated with nodule-related phenotypes in *Medicago truncatula*. We further investigated a subset of these genes and confirmed that several had existing evidence linking them nodulation, including MEDTR2G101090 (PEN3-like), a previously validated gene associated with nodule number.

## INTRODUCTION

The ability to convert atmospheric nitrogen into usable forms makes legumes an integral part of the plant ecosystem. Unfortunately, the expected increase in human population size over the next several decades will require a higher amount of nitrogen than current legume cropping systems can fulfill (Smil, 1999). This increase in demand requires that researchers better understand and improve nitrogen fixation in current legume species. One species in particular, Medicago truncatula, is widely considered a model species for understanding nitrogen fixation due to its diploid nature, seed to seed generation time, small genome size, and the vast amount of genomic resources (Young and Udvardi, 2009). Although previous studies have identified genes associated with nodulation (Oldroyd et al., 2001; Curtin et al., 2017; VandenBosch, 2003; Elise et al., 2005; Combier et al., 2006; Wasson, 2006), the trait is highly polygenic, and a large number of genes involved in nodulation remain to be discovered. One way researchers have tried to overcome this obstacle is through the use of Genome-wide association studies (GWAS).

Genetic analysis performed on standing collections of diverse lines or accessions reveals the locations of historical recombination that differentiate each genotype. GWAS leverage this information to discover associations between genetic markers and a phenotype of interest that exhibits variation within the population. However, these strong associations typically implicate genomic regions that are too large to allow for the identification of the specific gene that underlies this variation (Breseghello and Coelho, 2013; Flint-Garcia et al., 2005; Visscher et al., 2012). In most cases, further investigation is required to identify genes surrounding each marker that may be associated with the phenotype. Furthermore, it is possible that numerous markers truly associated with the trait are not identified as significant in GWAS, due to stringent statistical cutoffs (Storey and Tibshirani, 2003; Johnson et al., 2010; Sham and Purcell, 2014). Conversely, lowering the statistical threshold introduces false positives that are problematic for further analysis (Korte and Farlow, 2013).

Advances in next-generation sequencing technologies have allowed researchers to generate numerous reference genomes for a variety of plant species. However, many of the genes within these species remain functionally uncharacterized, limiting the amount of biological information available to interpret a candidate gene’s effect on a specific phenotype. Using technologies such as RNA-seq and microarrays, it is possible to measure quantitative levels of expression throughout the genome across multiple samples. Based on a collection of genome-wide expression profiles collected from various tissues, species, and/or environments, one can construct a co-expression network by measuring similarity between all pairs of genes’ expression profiles, where strongly connected edges indicate that two genes exhibit highly similar patterns of expression (Usadel et al., 2009; Stuart, 2003) (Aoki et al., 2007). These networks provide a powerful resource for understanding gene function, particularly for uncharacterized genes, as the data-derived relationships allow one to establish a functional context for a gene, even when formal annotations do not exist.

A recent study described a new framework to integrate co-expression networks with GWAS as a means to identify candidate genes (Schaefer et al., 2018). In maize, they ran several GWAS to identify a SNPs associated with elemental accumulation in seeds. Although they were able to identify significant markers associated with regions of the genome, in most cases, they were left with hundreds of markers per trait that often implicated linked genomic regions that could not be resolved to individual candidate genes. They further built three co-expression networks, two from publicly available data and one from root tissue designed to represent the phenotype measured in the respective GWAS. Schaefer et al. 2018 integrated the significant markers for each trait with the co-expression networks using their Camoco framework to identify and better prioritize candidate genes associated with elemental accumulation.

Here, we apply this framework to *Medicago truncatula* using publicly available expression datasets, and markers from a previously published GWAS focused on nodulation traits. We demonstrate that Camoco framework, originally established in maize, indeed generalizes to other species and traits, and provides an effective means of pinpointing candidate causal genes associated with nodulation.

## RESULTS AND DISCUSSION

### Integration of nodule focused genome-wide association study with co-expression networks

To identify candidate genes associated with nodulation traits, we used a previously published GWAS (Stanton-Geddes et al., 2013) as well as two publicly available RNA-seq datasets. The GWAS consisted of 226 *M. truncatula* accessions that were previously grown in replicate and phenotyped for five different nodulation traits as well as flowering time, trichrome density and height. By manually inspecting their most significant 50-200 SNPs ranked by p-value, the authors discovered several genes near significant SNPs that were previously associated with nodulation traits (Stanton-Geddes et al., 2013). Similar to other GWAS studies, the authors focused on genes that either contained or were directly adjacent to significant markers even though, in some cases, other genes may also be plausible candidates given their linkage to the significant markers (Branca et al., 2011). We selected a subset of these traits and markers from the study to serve as input for the GWAS/co-expression Camoco pipeline (Table S1).

As a basis for our co-expression networks, we used two publicly available RNA-seq expression data sets. The data consisted of 138 samples consisting of three different genotypes, three different tissues, four different rhizobium treatments, and presence-absence of nitrogen (Table S2). We then built six different co-expression networks using Camoco (https://github.com/schae234/Camoco) (Schaefer et al., 2018). Four of the six networks were constructed from a single tissue type (Leaf, Root, Nodule, JQL_Nodule), and the other two networks (referred to as the “General” network and “JQL” network) were constructed from a combination of different tissue types (Table S3). The diversity of tissue types within each co-expression network allows for the detection of signals corresponding to different biological processes that may have remained undiscovered if all samples were combined into one large network (Schaefer et al., 2014, 2018). The total number of genes that passed the co-expression network construction phase was relatively consistent among the four networks, with each network consisting of roughly 22,000 genes (Table S3). Genes that were excluded from each network were either not expressed, or did not exhibit enough variation in expression between samples to robustly measure covariation. The smaller number of genes within the nodule-specific network was expected, as fewer genes are expressed in nodule tissue relative to others (Benedito et al., 2008).

To test whether the networks were capturing biologically meaningful relationships, we measured the enrichment in each network for known biological relationships. Using sets of genes coannotated to the same Gene Ontology (GO) term, the relative density (how highly an established set of functionally related genes are co-expressed with each other) was measured and compared to density values of randomly sampled gene sets of the same size. All six networks demonstrated functional enrichment of at least ten-fold (Figure S1), indicating that many more GO terms exhibited evidence of co-expression than expected by chance for all six networks.

Using the six co-expression networks and selected GWAS markers, we applied the Camoco pipeline to prioritize candidate causal genes. Briefly, Camoco, which was originally described in Schaefer et al. 2018, evaluates candidate genes linked to significant GWAS markers on the basis of their co-expression with genes linked to other significant GWAS marker based on the assumption that some causal genes should exhibit strong co-expression relationships with other genes associated with the trait. Camoco is depicted in Figure 1, and the details of this analysis are provided in the Methods section. Any genes reported by Camoco with an FDR < 0.35 were considered candidate genes and included in further analysis.

**Figure 1.**
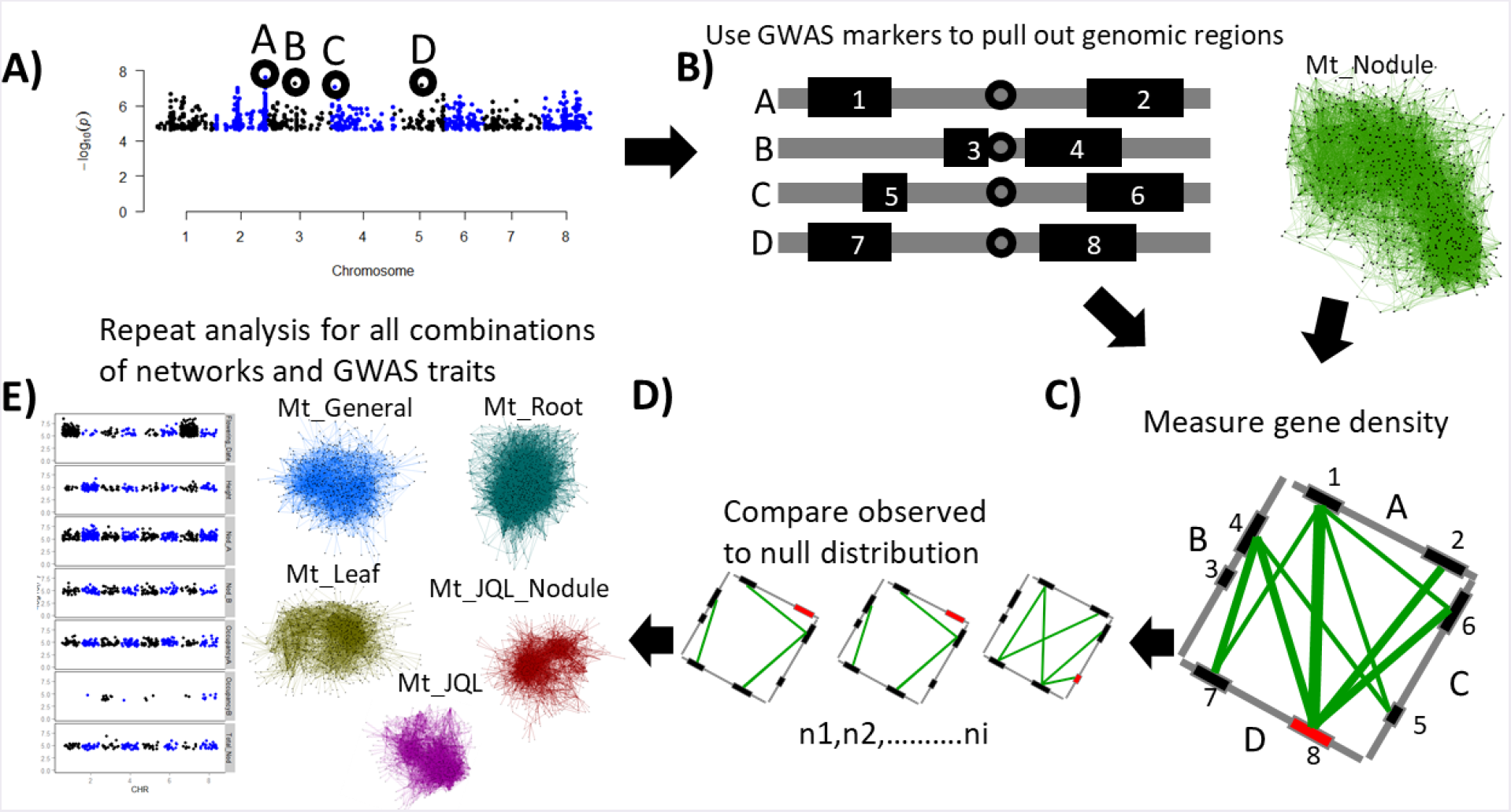
GWAS and co-expression pipeline. GWAS and co-expression pipeline using Camoco. A) Manhattan plot represents DNA markers used as input for Camoco, bold black circles represent a subset of markers used for illustrative purposes. B) Regions along a chromosome from previously selected markers are represented as grey bars, genes are represented as black rectangles. C) Genes from previously identified intervals are then selected from the co-expression network for per-gene network density measurements. Colored lines represent the strength of co-expression between two genes in a co-expression network. Wider lines, represent gene pairs that are more strongly co-expressed. The red box represents the current gene being measured for density. D) Per-gene density measurement of random sub-networks equal in size to the testing set. E) Other GWAS traits and networks used for analysis.

The results of the Camoco framework yielded 489 high-confidence candidate genes across all GWAS trait and network combinations. We also measured the number that persisted at more stringent FDR cutoffs, and indeed we were able to discover genes across a range of FDRs (FDR < 0.2: 172 genes; FDR < 0.1: 32 genes; FDR < 0.05: 3 genes). Analysis of the Nod_A trait (strain occupancy in the top 5 cm of roots) with the Mt_JQL_Nodule network combination, revealed a high amount of network connectivity between genes. To illustrate the basis for highly prioritized candidate genes, we highlight the observed co-expression relationships for MEDTR2G101090 (Figure 2), which was one of the top prioritized candidate genes for the Nod_A. MEDTR2G101090 is linked with a significant GWAS marker and is highly co-expressed with genes linked to significant loci on several other chromosomes (Figure 2), suggesting that the Camoco framework is discovering meaningful relationships.

**Figure 2.**
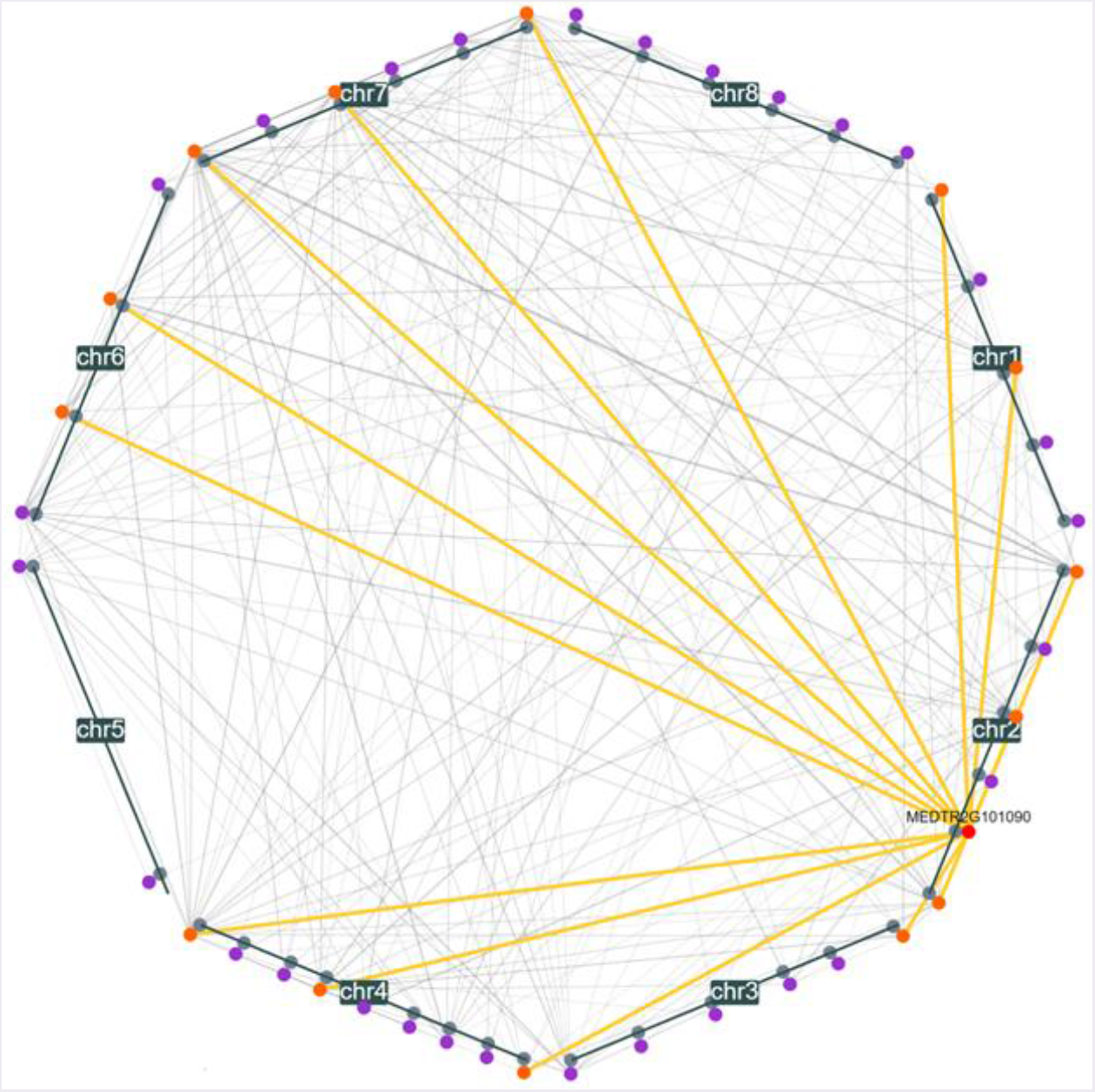
Nodule_A discoverable genes in the Mt_JQL_Nodule network. Chromosome-centric diagram of the connectivity of discoverable genes (FDR < 0.35), focused on coexpression neighbors of the candidate, MEDTR2G101090, within the JQL_nodule network for the Nod_A trait. Grey circles represent GWAS markers, colored circles represent genes, with MEDTR2G101090 in red, its first neighbors in orange, and other discoverable genes in purple. Grey lines represent co-expression between genes (minimum Z-Score of 2.5); the wider the line, the stronger the co-expression between genes.

### Importance of trait and tissue specificity in co-expression networks

The number of high-confidence candidate genes discovered by Camoco varied significantly across different combinations of traits, networks, and parameters (Figure 3). Interestingly, the nodule based Mt_JQL_Nodule co-expression network combined with the Nod_A trait yielded the most high-confidence candidate genes across all network-trait combinations, which likely reflects a strong match between the tissue in which expression covariation was measured and the biology of the phenotype of interest (in this case, both focused on nodules). Surprisingly, the root-based network performed the worst even though we expected strong biological relevance for nodulation based traits. It was the poorest performer across all GWAS traits, only producing a few candidate genes for the Nod_B trait (strain occupancy below the top 5 cm of roots). This result could possibly be due to the timepoint at which RNA was extracted from the roots. For example, if RNA was extracted at an earlier timepoint when nodules were still early in development, there may have been more informative expression patterns, allowing for the discovery of candidate genes.

**Figure 3.**
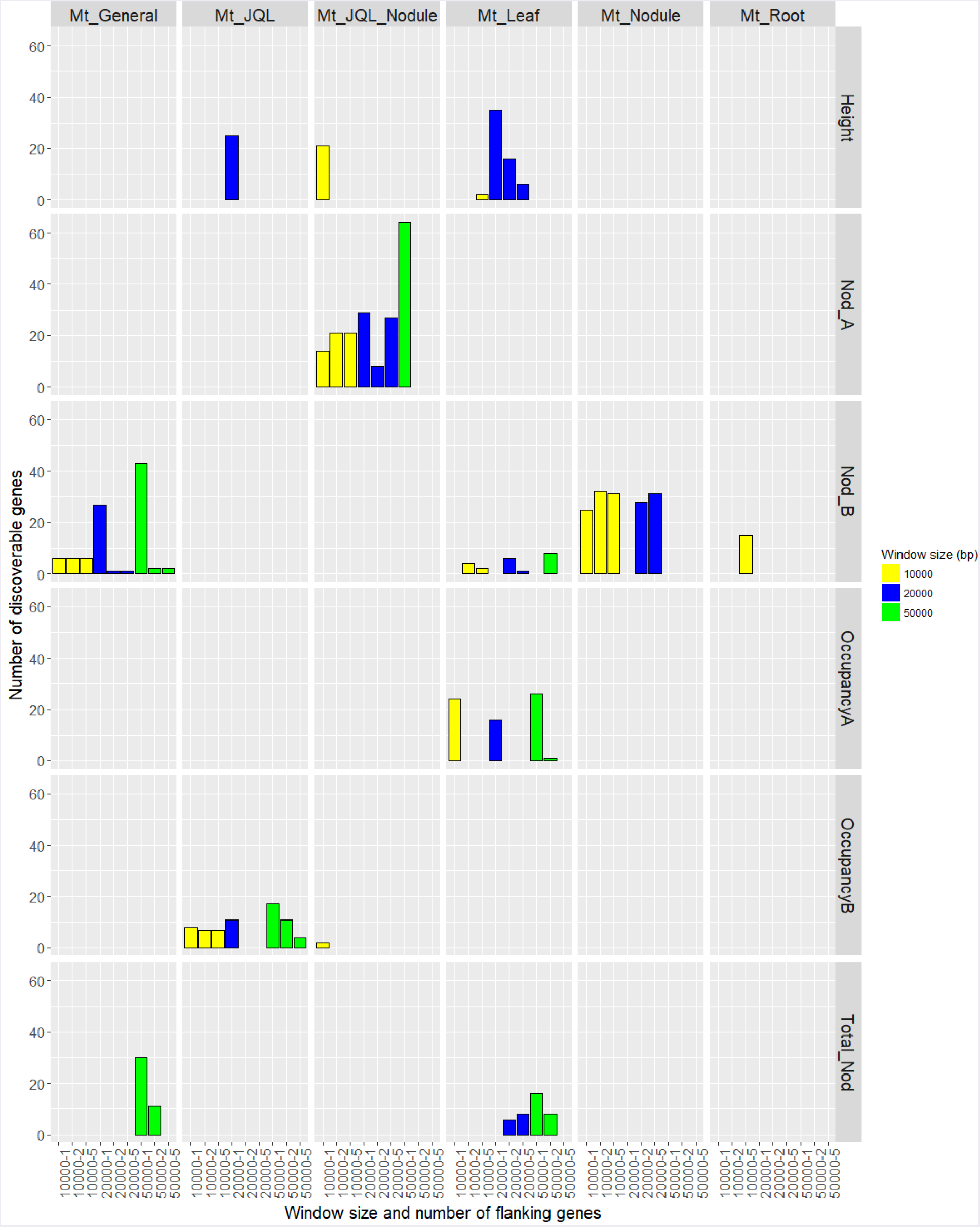
Co-expression/GWAS discoverable gene summary. Number of discoverable genes (FDR < 0.35) obtained from co-expression/GWAS integration. Colors represent the window size parameters used for our analysis.

The leaf network was the only network that consistently identified candidates for the height trait. While this is biologically unsurprising, this network also discovered significant genes for a few nodulation traits, suggesting that there are processes detectable in leaf tissue with relevance to nodulation. The General network, which consisted of the largest number of samples and tissue types only generated a few candidates for the Nod_B phenotype.

These results suggest that the context from which the co-expression network was derived, and its relation to the GWAS phenotype, play an important role in determining whether the Camoco approach is able to prioritize high-confidence genes. Notably, our results suggest that combining many different types of tissue into one large network does not perform well as a smaller, more concise, tissue-focused network, even though it is based on a larger set of expression profiles. One reason for this is that combining expression data from very different contexts introduces more variation across each gene’s profile, but that variation likely reveals generic modules that represent large sets of genes that simply are expressed in the same subsets of tissues. In contrast, networks derived from specific tissues capture more subtle covariation that reflects co-regulated genes functioning in processes relevant to that tissue that may otherwise be lost in larger sets of expression profiles.

### GWAS marker significance and proximity to genes are variable when integrating coexpression analysis

A common approach to interpreting GWAS studies is to manually inspect the most significant markers and look for candidates that are closest in proximity to the marker of interest. Unfortunately, the closest genes to GWAS markers may not always be the ones that are causally driving the association with the phenotype. When looking at the height trait in the leaf network, we see an increase in signal (i.e., number of Camoco-identified high-confidence candidate genes) as we increase the number of flanking genes surrounding each marker (Figure S2). When the window size is increased from 10 kb to 20kb, we see that the signal drastically increases, indicating that there are genes further out from the marker that are highly co-expressed with a subset of these genes. However, when an even larger 50kb window is used, no high-confidence genes are reported. The loss of signal at the largest interval (50kb) is expected as the number of potential candidate genes per locus increases sharply (the large majority of them being false positive as one considers candidates further from the locus peak). Ultimately, this large number of false candidate genes obscures the identification of co-expression relationships among true causal genes, and the approach no longer works. This analysis suggests that several of the GWAS loci implicated for these traits are likely driven by causal genes that are not directly adjacent to the GWAS peaks.

Similarly, the constraint of only focusing on the most significant markers (e.g. derived from extremely conservative significance cutoffs on the association test) leaves other candidates that are truly associated with the phenotype neglected. The Camoco framework can provide filtering of false positives at lower significance thresholds, by integrating information from the co-expression network. For instance, if we used the common GWAS p-value cutoff of 5 × 10^-8^ (Fadista et al., 2016; Barsh et al., 2012; Panagiotou and Ioannidis, 2012), this would result in two GWAS markers from the Nod_A phenotype, which does not provide enough context for an approach like Camoco to prioritize candidate genes. Instead, we applied a less conservative threshold (p-value < 3 x 10^-5^), which resulted in 292 SNPs, which was able to produce several high-confidence candidate causal genes, which would have otherwise been ignored (Table S1). In general, of course the number of markers produced at any confidence threshold will depend on the trait’s genetic architecture and the study design, but this analysis suggests that the Camoco approach can better produce candidate genes with less conservative thresholds on marker association.

### Identification of nodulation-related genes using co-expression and GWAS

To identify a small set of the most promising high confidence candidate genes for more investigation, we further narrowed candidate genes lists for the Nod_A trait by focusing on genes that were consistently discovered across different parameter settings. Using the JQL_Nodule network, we narrowed the candidate gene lists by limiting candidate genes to those that appeared in at least three out of the nine (10kb, 20kb, 50kb genome window size by 1,2,5 flanking gene) combinations of parameter settings; this process resulted in 25 genes for further investigation (Table1). When viewing the strength of co-expression between these 25 genes within the nodule network, it was observed that the majority of the genes were connected and formed a single module (Figure 4).

**Table 1.**
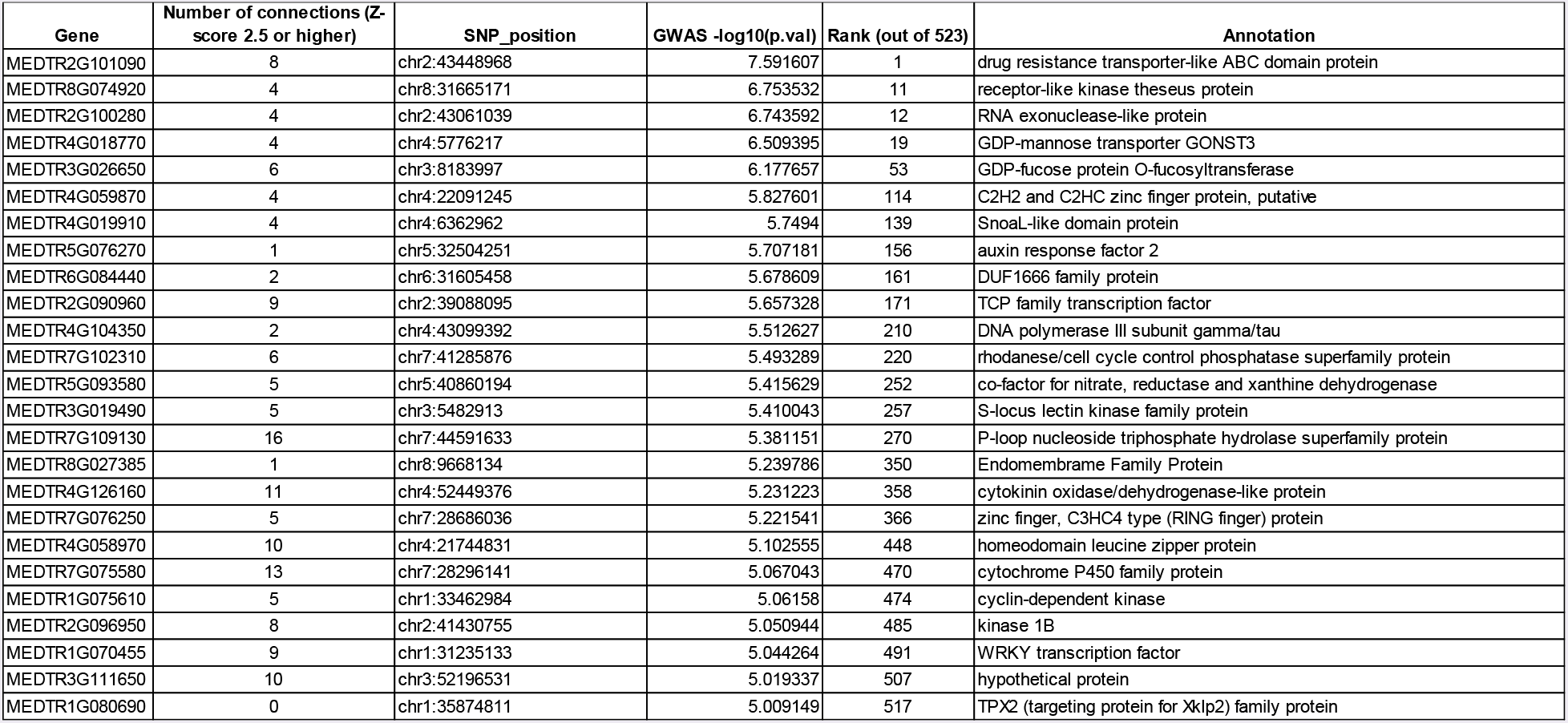
List of genes that were discoverable across all six parameters (10kb, 20kb and 1,2,5 flanking genes) for the Nod_A phenotype using the Mt_JQL Nodule GWAS.

**Figure 4.**
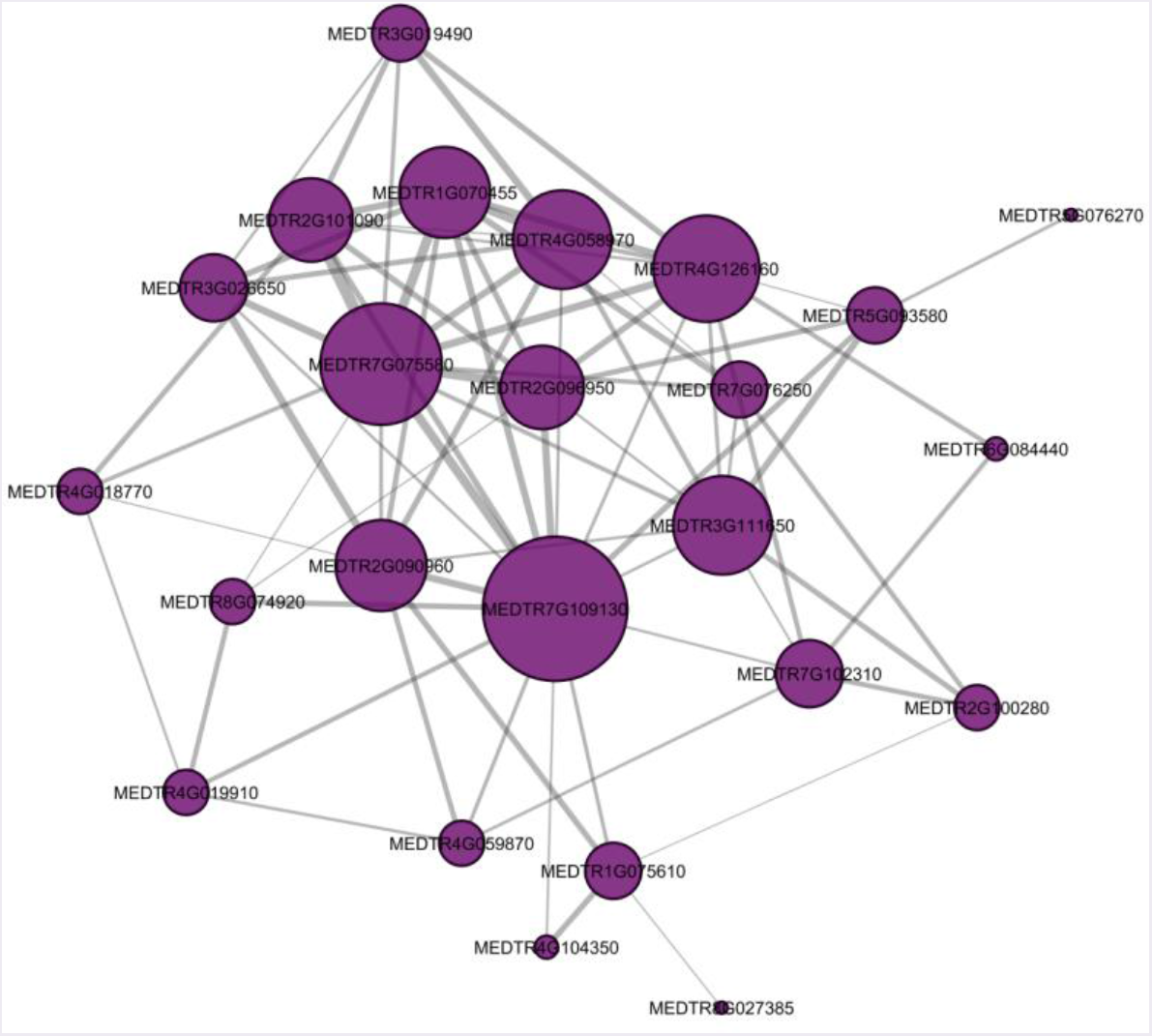
Overlap of Nod_A candidates in the Mt_JQL_Nodule network. Candidate genes for the JQL_nodule network for the Nod_A trait. Purple circles represent genes, and grey lines represent co-expression between genes (minimum Z-Score of 2.5). The larger the circle, the more connections it has with other genes. The wider the line, the stronger the co-expression between genes.

Interestingly, among those 25 candidate genes from the Nod_A analysis, was PEN3-like (MEDTR2G101090; Table 1), a gene that was associated with the most significant GWAS marker for the Nod_A trait (Stanton-Geddes et al., 2013). Functional validation of PEN3-like using CRISPR and Tnt1-mutated plants previously confirmed that loss-of-function of this gene resulted in decreased nodule number (Curtin et al., 2017). Another strong candidate among these 25 within the module was the hub gene (gene with the highest number of connections), MEDTR7G109130, which is annotated as a P-loop nucleoside triphosphate hydrolase superfamily protein and is known to play a role in nodulation (Jayaraman et al., 2017).

Because multiple co-expression networks were able to support the discovery of strong candidate genes for Nod_B, we defined a short list of high-confidence candidates by requiring high confidence genes to be consistently prioritized as high-confidence candidates across all networks for the Nod_B trait and were discovered across 4 or more parameter settings (Table 2). One promising gene, MEDTR1G012530, appeared as a candidate for 9 out of the 20 parameter settings that resulted in at least one candidate gene discovery. This gene is annotated as a TPX2 (targeting protein for Xklp2) family protein and has been shown to be highly expressed during nodule formation (Jardinaud et al., 2016). Another promising candidate, MEDTR4G073400, which also appeared as candidate 9 times, is annotated as Synaptotagmins-1-related, which play a role in the formation of root nodules (Gavrin et al., 2017).

**Table 2:**
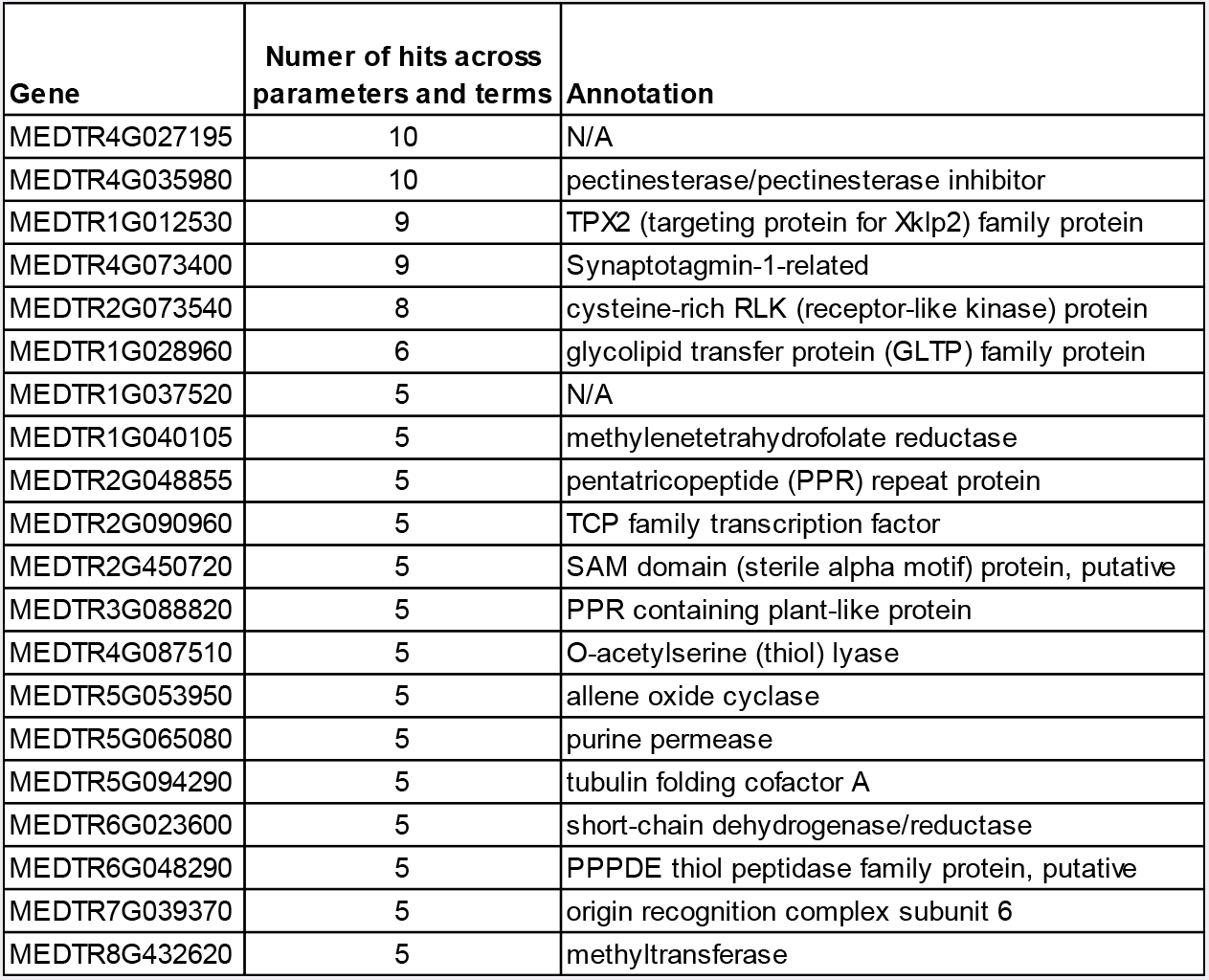
List of genes that were discoverable for at least 5 different parameters across all networks for the Nod_B trait

| Gene | Numer of hits across parameters and terms | Annotation |
| --- | --- | --- |
| MEDTR4G027195 | 10 | N/A |
| MEDTR4G035980 | 10 | pectinesterase/pectinesterase inhibitor |
| MEDTR1G012530 | 9 | TPX2 (targeting protein for Xklp2) family protein |
| MEDTR4G073400 | 9 | Synaptotagmin-1-related |
| MEDTR2G073540 | 8 | cysteine-rich RLK (receptor-like kinase) protein |
| MEDTR1G028960 | 6 | glycolipid transfer protein (GLTP) family protein |
| MEDTR1G037520 | 5 | N/A |
| MEDTR1G040105 | 5 | methylenetetrahydrofolate reductase |
| MEDTR2G048855 | 5 | pentatricopeptide (PPR) repeat protein |
| MEDTR2G090960 | 5 | TCP family transcription factor |
| MEDTR2G450720 | 5 | SAM domain (sterile alpha motif) protein, putative |
| MEDTR3G088820 | 5 | PPR containing plant-like protein |
| MEDTR4G087510 | 5 | O-acetylserine (thiol) lyase |
| MEDTR5G053950 | 5 | allene oxide cyclase |
| MEDTR5G065080 | 5 | purine permease |
| MEDTR5G094290 | 5 | tubulin folding cofactor A |
| MEDTR6G023600 | 5 | short-chain dehydrogenase/reductase |
| MEDTR6G048290 | 5 | PPPDE thiol peptidase family protein, putative |
| MEDTR7G039370 | 5 | origin recognition complex subunit 6 |
| MEDTR8G432620 | 5 | methyltransferase |

Overall, these results demonstrate that the integration of co-expression networks to interpret GWAS results was able to effectively prioritize genes causally associated with nodulation processes. The genes that are directly connected to PEN3-like would serve as valuable candidates for follow-up studies due to their similarity in expression profiles across tissues. Another approach to prioritizing candidates from among the set produced by the Camoco analysis is to rank based on their linked GWAS marker’s significance value. For instance, the candidate causal gene associated with the marker with the highest significance was the PEN3-like gene while our P-loop nucleoside triphosphate hydrolase superfamily protein hub gene was ranked 270 out of 523 significant markers input into the Camoco analysis.

## Conclusions

Using an *M. truncatula* GWAS focused on nodulation traits as well as expression data from different tissues, rhizobium strains, nitrogen treatments and accessions, we were able to identify a subset of genes surrounding GWAS markers that are highly co-expressed with one another. From these lists, we discovered a previously validated nodulation gene PEN3-like as well as several other genes whose annotations are associated with nodulation. Uncharacterized genes within our high-confidence lists are worthy of more in-depth follow-up studies using Tnt1 or CRISPR knockouts.

Schaefer et al. 2018 developed the Camoco framework and integrated co-expression networks and GWAS in maize in order to capture variation associated with elemental uptake in seeds. Our current study used a higher-density GWAS that focused on a different phenotype, different plant species, and an expression data set that was not explicitly created for this study. One common theme between the studies is that the choice of the co-expression network matters; specifically, tissue-relevant networks derived from expression variation across diverse genotypes appear to perform the best in ranking candidate genes. This was true in maize, and we report here that this is also true in Medicago. We believe this result is likely to generalize to many other contexts, and it suggests as a community, more emphasis in the generation genotype-focused networks would be worthwhile if we hope to build resources for functional interpretation of phenotype-associated variants. It is also important to mention that we were able to generate a panel of high confidence candidate genes using two independent datasets that were not generated specifically for this study.

The majority of candidate genes discovered in this analysis would have mostly likely been neglected by traditional GWAS analyses unless they were under the most significant markers. By combining co-expression networks with GWAS, the functional relationship between genes related to the GWAS phenotype are more likely to be discovered. It is also important to note that based on our analysis, in many cases, the nearest gene to a marker was not the gene predicted to be causally associated with the phenotype.

In general, we demonstrate that the Camoco framework for integrating co-expression networks with GWAS generalizes beyond the species for which it was originally developed and applied (maize ionomic traits), as it shows utility for prioritizing genes related to nodulation in *Medicago truncatula*. Based on these results, we expect that the approach will generalize to a wide variety of other species and traits as well.

## Acknowledgments

We would like to thank the University of Minnesota Office of Information Technology for accommodating our data storage needs and the Department of Computer Science at the University of Minnesota for server maintenance and support. This work was supported by funding from the National Science Foundation (IOS-1237993) with partial funding from (IOS-1126950). Funding sources played no role in the design of this study or the collection, analysis, and the interpretation of data and in writing the manuscript.

## MATERIAL AND METHODS

### *Medicago* experimental design and sample extraction

Three accessions from the Medicago HapMap project (HM56, HM101, HM340) were grown in greenhouse conditions. Rhizobium strains S. meliloti (KH46c) and S. medicae (WSM419), as well as nitrogen, were applied to the soil shortly after planting. Tissues were harvested and frozen in liquid nitrogen 31 days after planting. RNA was extracted using the Qiagen RNeasy Plant mini kit (Product ID: 74903). Individual nodules were pooled and extracted as a single sample for each plant.

### Generation of expression data

RNA from 138 samples were sequenced by the University of Minnesota’s Genomic Center using Illumina HiSeq2500 100bp single-end reads. One sample required resequencing (L88), which resulted in 125bp reads. Samples were barcoded and multiplexed using Illumina TruSeq HT adapters. Fastq files were checked with Fastqc version 0.11.5 and adapters were trimmed using cutadapt version 1.8.1 with non-default parameters -m 40 and -q 30 (Andrews, 2010; Martin, 2011). Reads were then aligned to Mt_4.0 gene models, and reference (http://jcvi.org/medicago/) using STAR 2.5.3a (Dobin et al., 2013), then filtered based on unique mapping scores, sorted and indexed using samtools version 1.6 (Li et al., 2009). FPKM values were generated using Cufflinks version 2.2.1 using non-default parameters of -I 20000 and --min-intron-length 5. Raw sequencing files are publicly available on the NCBI SRA (PRJNA327225 and PRJNA449544).

### Co-expression network construction and genome-wide association study integration

Methods used were similar to those in the previously mentioned co-expression GWAS integration study (Schaefer 2017). Briefly, Camoco takes a set of SNP’s as input and uses their location within a genome as well the number of genes flanking a marker within a given window size to extract genes lists for testing (Figure 1). If there are multiple significant SNPs appearing within the same window, then all but the most significant SNP is discarded (Table S1). Once genes are selected for testing; each gene is then measured to see how well it is co-expressed with other genes also linked to the significant markers associated with the trait of interest. Once a network statistic (either density or locality, see Schaefer 2017) is generated, Camoco will resample (1000 times) a random set of genes equal in size to the test set to establish a null distribution for estimating significance of the observed statistic. To account for the varying amount of linkage disequilibrium across the genome, we used 10kb, 20kb and 50kb window sizes and 1, 2, and 5 flanking genes (Stanton-Geddes et al., 2013). Any gene that had an FDR < 0.35 was called “candidate” and included in further analysis.

FPKM expression tables were used as input into Camoco (https://github.com/schae234/Camoco) using the Mt_4.1 reference genome. Non-default parameters used to build each network included rawtype=‘RNASEQ’, max_gene_missing_data=0.5, max_accession_missing_data=0.5, min_single_sample_expr=1, min_expr=0.001, quantile=False, max_val=300, sep=‘,’. Network health statistics were generated using GO terms from (http://jcvi.org/medicago) and 1000 bootstraps. SNPs were integrated into Camoco using built-in functions, and per gene, density measurements were run with 1,000 bootstraps. Figures were created using ggplot2 (Wickham, 2006).

---

**Figure S1.**
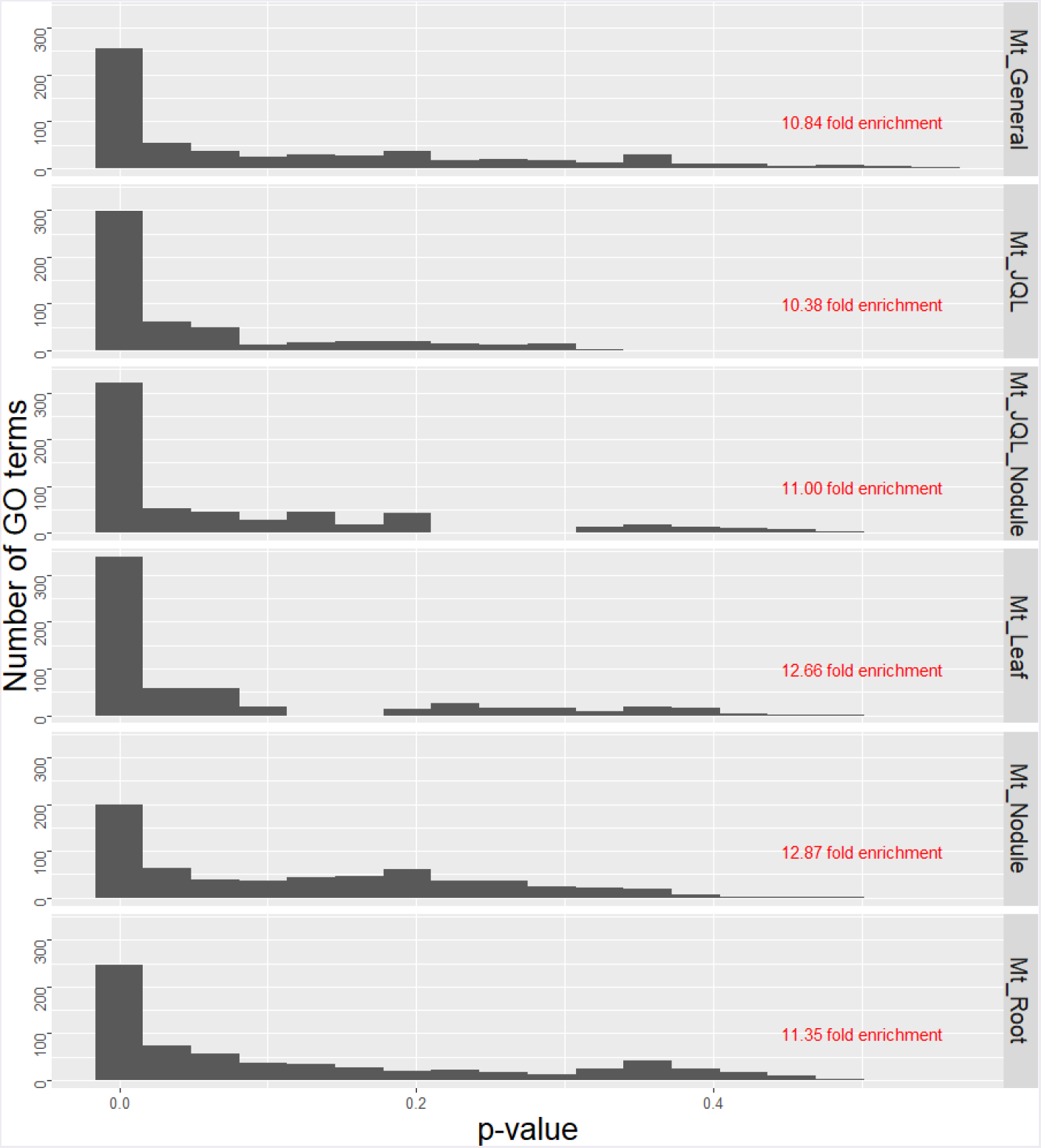
Network GO term enrichment. Distribution of p-values from density-based GO-term enrichment. A histogram of p-values for each density-based GO-term enrichment test based on its density, relative to the distribution of density values from random gene sets similar in size.

**Figure S2.**
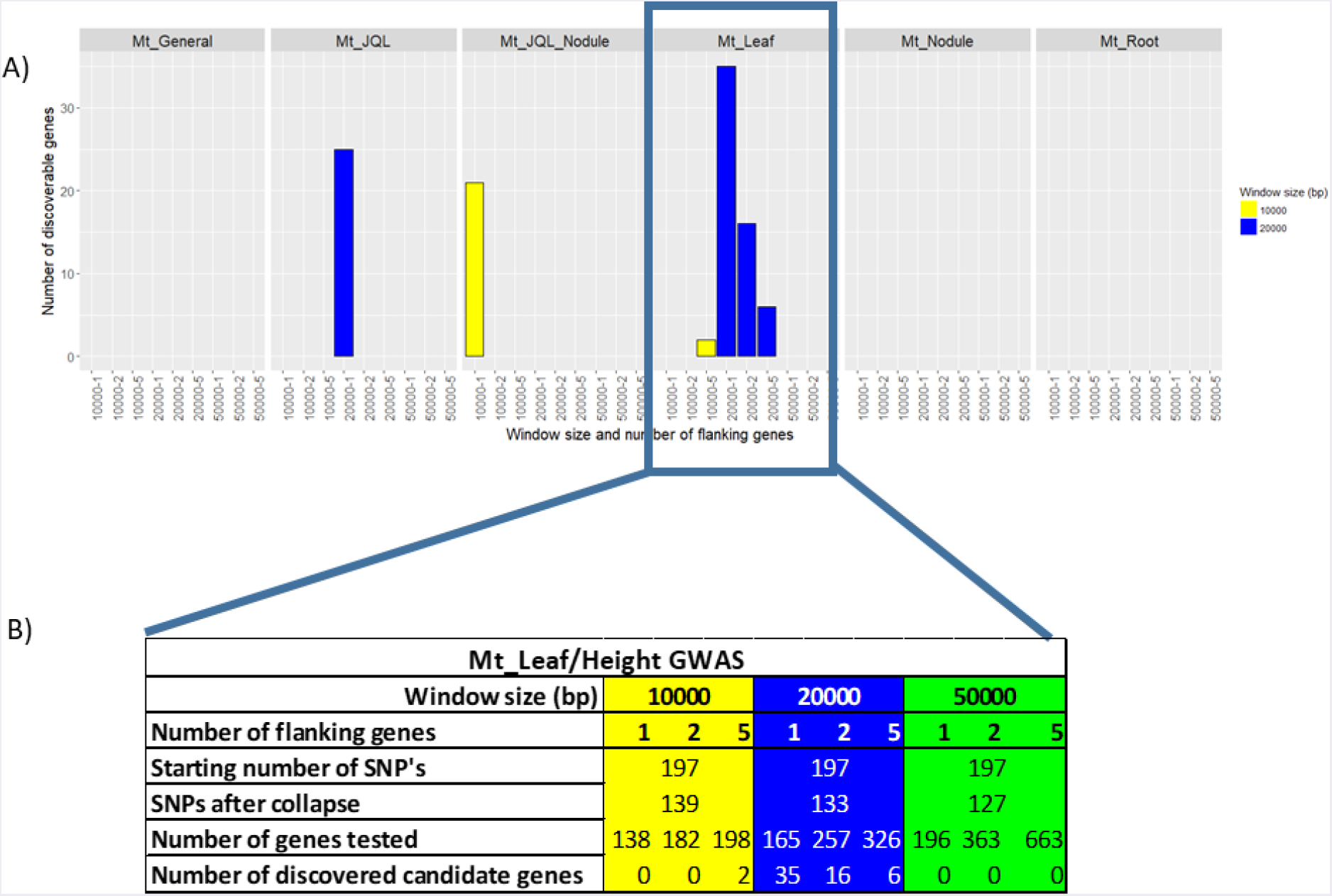
Co-expression/Height GWAS discoverable gene summary. Flow chart of candidate gene identification in the height GWAS trait. A) Number of discoverable genes (FDR < 0.35) using the height GWAS with each co-expression network. Colors represent the window size parameter use with Camoco. B) The number of SNP’s and genes that were included in each analysis.

**Table S1.** GWAS trait information and the number of SNP’s used for analysis. “Collapse” refers to SNP’s removed due to overlapping windows between sets of SNPs

| GWAS Trait | Description | Number of SNP's | SNP's included after collapse (window size) |  |  | Min p-value |
| --- | --- | --- | --- | --- | --- | --- |
|  |  |  | 10kb | 20kb | 50kb |  |
| Height | Plant height | 197 | 139 | 133 | 127 | 3.00E-05 |
| Total_Nod | Total number of nodules | 163 | 124 | 122 | 119 | 3.00E-05 |
| Nod_A | Total number of nodules in the top 5 cm of roots | 523 | 294 | 275 | 255 | 9.96E-06 |
| Nod_B | Total number of nodules below the top 5 cm of roots | 232 | 185 | 178 | 165 | 3.00E-05 |
| Flowering_Date | Flowering date | 550 | 150 | 120 | 100 | 6.94E-06 |
| OccupancyA | Strain occupancy in the top 5 cm of roots | 292 | 230 | 226 | 209 | 3.00E-05 |
| OccupancyB | Strain occupancy below the top 5 cm of roots | 27 | 17 | 17 | 14 | 9.61E-05 |

**Table S2.** Metadata regarding the 138 samples used for analysis

| Sample ID | Tissue | M. truncatula accession | Sinorhizobium species and strain | Nitrogen | D7 Index | Barcode | D5 Index | Barcode |
| --- | --- | --- | --- | --- | --- | --- | --- | --- |
| N128 | Nodule | HM056 | S. meliloti (KH46c) | 0 | D701 | ATTACTCG | D501 | TATAGCCT |
| N86 | Nodule | HM056 | S. meliloti (KH46c) | 0 | D702 | TCCGGAGA | D501 | TATAGCCT |
| N73 | Nodule | HM056 | S. medicae (WSM419) | 0 | D704 | GAGATTCC | D501 | TATAGCCT |
| N137 | Nodule | HM056 | S. medicae (WSM419) | 0 | D705 | ATTCAGAA | D501 | TATAGCCT |
| N48 | Nodule | HM056 | S. medicae (WSM419) | 0 | D706 | GAATTCGT | D501 | TATAGCCT |
| N88 | Nodule | HM101 | Both | 0 | D707 | CTGAAGCT | D501 | TATAGCCT |
| N103 | Nodule | HM101 | Both | 0 | D708 | TAATGCGC | D501 | TATAGCCT |
| N25 | Nodule | HM101 | Both | 0 | D709 | CGGCTATG | D501 | TATAGCCT |
| N9 | Nodule | HM101 | S. meliloti (KH46c) | 0 | D710 | TCCGCGAA | D501 | TATAGCCT |
| N121 | Nodule | HM101 | S. meliloti (KH46c) | 0 | D711 | TCTCGCGC | D501 | TATAGCCT |
| N39 | Nodule | HM101 | S. meliloti (KH46c) | 1 | D712 | AGCGATA<br>G | D501 | TATAGCCT |
| N75 | Nodule | HM101 | S. meliloti (KH46c) | 1 | D701 | ATTACTCG | D502 | ATAGAGGC |
| N146 | Nodule | HM101 | S. meliloti (KH46c) | 1 | D702 | TCCGGAGA | D502 | ATAGAGGC |
| N83 | Nodule | HM101 | S. medicae (WSM419) | 0 | D704 | GAGATTCC | D502 | ATAGAGGC |
| N56 | Nodule | HM101 | S. medicae (WSM419) | 0 | D705 | ATTCAGAA | D502 | ATAGAGGC |
| N14 | Nodule | HM101 | S. medicae (WSM419) | 0 | D706 | GAATTCGT | D502 | ATAGAGGC |
| N64 | Nodule | HM101 | S. medicae (WSM419) | 0 | D707 | CTGAAGCT | D502 | ATAGAGGC |
| N122 | Nodule | HM101 | S. medicae (WSM419) | 1 | D708 | TAATGCGC | D502 | ATAGAGGC |
| N46 | Nodule | HM101 | S. medicae (WSM419) | 1 | D709 | CGGCTATG | D502 | ATAGAGGC |
| N41 | Nodule | HM101 | S. medicae (WSM419) | 1 | D710 | TCCGCGAA | D502 | ATAGAGGC |
| N107 | Nodule | HM101 | S. medicae (WSM419) | 1 | D711 | TCTCGCGC | D502 | ATAGAGGC |
| N62 | Nodule | HM340 | Both | 0 | D712 | AGCGATA<br>G | D502 | ATAGAGGC |
| N21 | Nodule | HM340 | Both | 0 | D701 | ATTACTCG | D503 | CCTATCCT |
| N160 | Nodule | HM340 | S. meliloti (KH46c) | 0 | D702 | TCCGGAGA | D503 | CCTATCCT |
| N115 | Nodule | HM340 | S. meliloti (KH46c) | 1 | D704 | GAGATTCC | D503 | CCTATCCT |
| N131 | Nodule | HM340 | S. meliloti (KH46c) | 1 | D705 | ATTCAGAA | D503 | CCTATCCT |
| N143 | Nodule | HM340 | S. meliloti (KH46c) | 1 | D706 | GAATTCGT | D503 | CCTATCCT |
| N80 | Nodule | HM340 | S. medicae (WSM419) | 0 | D707 | CTGAAGCT | D503 | CCTATCCT |
| N92 | Nodule | HM340 | S. medicae (WSM419) | 0 | D708 | TAATGCGC | D503 | CCTATCCT |
| N26 | Nodule | HM340 | S. medicae (WSM419) | 0 | D709 | CGGCTATG | D503 | CCTATCCT |
| N8 | Nodule | HM340 | S. medicae (WSM419) | 0 | D710 | TCCGCGAA | D503 | CCTATCCT |
| N42 | Nodule | HM340 | S. medicae (WSM419) | 1 | D711 | TCTCGCGC | D503 | CCTATCCT |
| N47 | Nodule | HM340 | S. medicae (WSM419) | 1 | D712 | AGCGATA<br>G | D503 | CCTATCCT |
| N111 | Nodule | HM340 | S. medicae (WSM419) | 1 | D701 | ATTACTCG | D504 | GGCTCTGA |
| N120 | Nodule | HM056 | S. medicae (WSM419) | 0 | D704 | GAGATTCC | D502 | ATAGAGGC |
| N40 | Nodule | HM101 | S. meliloti (KH46c) | 1 | D705 | ATTCAGAA | D502 | ATAGAGGC |
| N11 | Nodule | HM340 | S. meliloti (KH46c) | 0 | D706 | GAATTCGT | D502 | ATAGAGGC |
| R51 | Root | HM101 | S. meliloti (KH46c) | 0 | D702 | TCCGGAGA | D504 | GGCTCTGA |
| R5 | Root | HM101 | S. meliloti (KH46c) | 0 | D705 | ATTCAGAA | D504 | GGCTCTGA |
| R125 | Root | HM101 | None | 1 | D706 | GAATTCGT | D504 | GGCTCTGA |
| R171 | Root | HM101 | None | 1 | D707 | CTGAAGCT | D504 | GGCTCTGA |
| R142 | Root | HM101 | None | 1 | D708 | TAATGCGC | D504 | GGCTCTGA |
| R83 | Root | HM101 | S. medicae (WSM419) | 0 | D709 | CGGCTATG | D504 | GGCTCTGA |
| R56 | Root | HM101 | S. medicae (WSM419) | 0 | D710 | TCCGCGAA | D504 | GGCTCTGA |
| R14 | Root | HM101 | S. medicae (WSM419) | 0 | D711 | TCTCGCGC | D504 | GGCTCTGA |
| R64 | Root | HM101 | S. medicae (WSM419) | 0 | D712 | AGCGATA<br>G | D504 | GGCTCTGA |
| R160 | Root | HM340 | S. meliloti (KH46c) | 0 | D701 | ATTACTCG | D505 | AGGCGAAG |
| R11 | Root | HM340 | S. meliloti (KH46c) | 0 | D702 | TCCGGAGA | D505 | AGGCGAAG |
| R44 | Root | HM340 | None | 1 | D704 | GAGATTCC | D505 | AGGCGAAG |
| R13 | Root | HM340 | None | 1 | D705 | ATTCAGAA | D505 | AGGCGAAG |
| R33 | Root | HM340 | None | 1 | D706 | GAATTCGT | D505 | AGGCGAAG |
| R80 | Root | HM340 | S. medicae (WSM419) | 0 | D707 | CTGAAGCT | D505 | AGGCGAAG |
| R92 | Root | HM340 | S. medicae (WSM419) | 0 | D708 | TAATGCGC | D505 | AGGCGAAG |
| R26 | Root | HM340 | S. medicae (WSM419) | 0 | D709 | CGGCTATG | D505 | AGGCGAAG |
| R8 | Root | HM340 | S. medicae (WSM419) | 0 | D710 | TCCGCGAA | D505 | AGGCGAAG |
| R9 | Root | HM101 | S. meliloti (KH46c) | 0 | D707 | CTGAAGCT | D502 | ATAGAGGC |
| R34 | Root | HM340 | S. meliloti (KH46c) | 0 | D708 | TAATGCGC | D502 | ATAGAGGC |
| L70 | Leaf | HM056 | S. meliloti (KH46c) | 0 | D712 | AGCGATA<br>G | D505 | AGGCGAAG |
| L128 | Leaf | HM056 | S. meliloti (KH46c) | 0 | D701 | ATTACTCG | D506 | TAATCTTA |
| L152 | Leaf | HM056 | S. meliloti (KH46c) | 0 | D702 | TCCGGAGA | D506 | TAATCTTA |
| L86 | Leaf | HM056 | S. meliloti (KH46c) | 0 | D709 | CGGCTATG | D502 | ATAGAGGC |
| L20 | Leaf | HM056 | None | 1 | D704 | GAGATTCC | D506 | TAATCTTA |
| L59 | Leaf | HM056 | None | 1 | D705 | ATTCAGAA | D506 | TAATCTTA |
| L60 | Leaf | HM056 | None | 1 | D706 | GAATTCGT | D506 | TAATCTTA |
| L61 | Leaf | HM056 | None | 1 | D707 | CTGAAGCT | D506 | TAATCTTA |
| L120 | Leaf | HM056 | S. medicae (WSM419) | 0 | D708 | TAATGCGC | D506 | TAATCTTA |
| L73 | Leaf | HM056 | S. medicae (WSM419) | 0 | D709 | CGGCTATG | D506 | TAATCTTA |
| L137 | Leaf | HM056 | S. medicae (WSM419) | 0 | D710 | TCCGCGAA | D506 | TAATCTTA |
| L48 | Leaf | HM056 | S. medicae (WSM419) | 0 | D711 | TCTCGCGC | D506 | TAATCTTA |
| L88 | Leaf | HM101 | Both | 0 | D712 | AGCGATA<br>G | D506 | TAATCTTA |
| L103 | Leaf | HM101 | Both | 0 | D701 | ATTACTCG | D507 | CAGGACGT |
| L25 | Leaf | HM101 | Both | 0 | D702 | TCCGGAGA | D507 | CAGGACGT |
| L158 | Leaf | HM101 | Both | 0 | D710 | TCCGCGAA | D502 | ATAGAGGC |
| L51 | Leaf | HM101 | S. meliloti (KH46c) | 0 | D704 | GAGATTCC | D507 | CAGGACGT |
| L9 | Leaf | HM101 | S. meliloti (KH46c) | 0 | D705 | ATTCAGAA | D507 | CAGGACGT |
| L5 | Leaf | HM101 | S. meliloti (KH46c) | 0 | D707 | CTGAAGCT | D507 | CAGGACGT |
| L39 | Leaf | HM101 | S. meliloti (KH46c) | 1 | D708 | TAATGCGC | D507 | CAGGACGT |
| L75 | Leaf | HM101 | S. meliloti (KH46c) | 1 | D709 | CGGCTATG | D507 | CAGGACGT |
| L146 | Leaf | HM101 | S. meliloti (KH46c) | 1 | D710 | TCCGCGAA | D507 | CAGGACGT |
| L40 | Leaf | HM101 | S. meliloti (KH46c) | 1 | D711 | TCTCGCGC | D507 | CAGGACGT |
| L125 | Leaf | HM101 | None | 1 | D712 | AGCGATAG | D507 | CAGGACGT |
| L171 | Leaf | HM101 | None | 1 | D701 | ATTACTCG | D508 | GTACTGAC |
| L142 | Leaf | HM101 | None | 1 | D702 | TCCGGAGA | D508 | GTACTGAC |
| L83 | Leaf | HM101 | S. medicae (WSM419) | 0 | D711 | TCTCGCGC | D502 | ATAGAGGC |
| L56 | Leaf | HM101 | S. medicae (WSM419) | 0 | D704 | GAGATTCC | D508 | GTACTGAC |
| L14 | Leaf | HM101 | S. medicae (WSM419) | 0 | D705 | ATTCAGAA | D508 | GTACTGAC |
| L64 | Leaf | HM101 | S. medicae (WSM419) | 0 | D706 | GAATTCGT | D508 | GTACTGAC |
| L62 | Leaf | HM340 | Both | 0 | D707 | CTGAAGCT | D508 | GTACTGAC |
| L21 | Leaf | HM340 | Both | 0 | D708 | TAATGCGC | D508 | GTACTGAC |
| L118 | Leaf | HM340 | Both | 0 | D709 | CGGCTATG | D508 | GTACTGAC |
| L49 | Leaf | HM340 | Both | 0 | D710 | TCCGCGAA | D508 | GTACTGAC |
| L160 | Leaf | HM340 | S. meliloti (KH46c) | 0 | D711 | TCTCGCGC | D508 | GTACTGAC |
| L11 | Leaf | HM340 | S. meliloti (KH46c) | 0 | D701 | ATTACTCG | D501 | TATAGCCT |
| L34 | Leaf | HM340 | S. meliloti (KH46c) | 0 | D702 | TCCGGAGA | D501 | TATAGCCT |
| L44 | Leaf | HM340 | None | 1 | D711 | TCTCGCGC | D501 | TATAGCCT |
| L13 | Leaf | HM340 | None | 1 | D704 | GAGATTCC | D501 | TATAGCCT |
| L33 | Leaf | HM340 | None | 1 | D705 | ATTCAGAA | D501 | TATAGCCT |
| L80 | Leaf | HM340 | S. medicae (WSM419) | 0 | D706 | GAATTCGT | D501 | TATAGCCT |
| L92 | Leaf | HM340 | S. medicae (WSM419) | 0 | D707 | CTGAAGCT | D501 | TATAGCCT |
| L26 | Leaf | HM340 | S. medicae (WSM419) | 0 | D708 | TAATGCGC | D501 | TATAGCCT |
| L8 | Leaf | HM340 | S. medicae (WSM419) | 0 | D709 | CGGCTATG | D501 | TATAGCCT |
| L121 | Leaf | HM101 | S. meliloti (KH46c) | 0 | D706 | GAATTCGT | D507 | CAGGACGT |
| JQL01 | Nodule | HM101 | S. meliloti (KH46c) | 0 | NA | NA | NA | NA |
| JQL02 | Nodule | HM101 | S. meliloti (KH46c) | 0 | NA | NA | NA | NA |
| JQL03 | Nodule | HM101 | S. meliloti (KH46c) | 0 | NA | NA | NA | NA |
| JQL04 | Nodule | HM101 | S. medicae (WSM419) | 0 | NA | NA | NA | NA |
| JQL05 | Nodule | HM101 | S. medicae (WSM419) | 0 | NA | NA | NA | NA |
| JQL06 | Nodule | HM101 | S. medicae (WSM419) | 0 | NA | NA | NA | NA |
| JQL07 | Root | HM101 | None | 0 | NA | NA | NA | NA |
| JQL08 | Root | HM101 | None | 0 | NA | NA | NA | NA |
| JQL09 | Root | HM101 | None | 0 | NA | NA | NA | NA |
| JQL10 | Nodule | HM056 | S. meliloti (KH46c) | 0 | NA | NA | NA | NA |
| JQL11 | Nodule | HM056 | S. meliloti (KH46c) | 0 | NA | NA | NA | NA |
| JQL12 | Nodule | HM056 | S. meliloti (KH46c) | 0 | NA | NA | NA | NA |
| JQL13 | Nodule | HM056 | S. medicae (WSM419) | 0 | NA | NA | NA | NA |
| JQL14 | Nodule | HM056 | S. medicae (WSM419) | 0 | NA | NA | NA | NA |
| JQL15 | Nodule | HM056 | S. medicae (WSM419) | 0 | NA | NA | NA | NA |
| JQL16 | Root | HM056 | None | 0 | NA | NA | NA | NA |
| JQL17 | Root | HM056 | None | 0 | NA | NA | NA | NA |
| JQL18 | Root | HM056 | None | 0 | NA | NA | NA | NA |
| JQL19 | Nodule | HM340 | *S. meliloti* (KH46c) | 0 | NA | NA | NA | NA |
| JQL20 | Nodule | HM340 | *S. meliloti* (KH46c) | 0 | NA | NA | NA | NA |
| JQL21 | Nodule | HM340 | *S. meliloti* (KH46c) | 0 | NA | NA | NA | NA |
| JQL22 | Nodule | HM340 | *S. medicae* (WSM419) | 0 | NA | NA | NA | NA |
| JQL23 | Nodule | HM340 | *S. medicae* (WSM419) | 0 | NA | NA | NA | NA |
| JQL24 | Nodule | HM340 | *S. medicae* (WSM419) | 0 | NA | NA | NA | NA |
| JQL25 | Root | HM340 | None | 0 | NA | NA | NA | NA |
| JQL26 | Root | HM340 | None | 0 | NA | NA | NA | NA |
| JQL27 | Root | HM340 | None | 0 | NA | NA | NA | NA |
| JQL28 | Nodule | HM034 | *S. meliloti* (KH46c) | 0 | NA | NA | NA | NA |
| JQL29 | Nodule | HM034 | *S. meliloti* (KH46c) | 0 | NA | NA | NA | NA |
| JQL30 | Nodule | HM034 | *S. meliloti* (KH46c) | 0 | NA | NA | NA | NA |
| JQL31 | Nodule | HM034 | *S. medicae* (WSM419) | 0 | NA | NA | NA | NA |
| JQL32 | Nodule | HM034 | *S. medicae* (WSM419) | 0 | NA | NA | NA | NA |
| JQL33 | Nodule | HM034 | *S. medicae* (WSM419) | 0 | NA | NA | NA | NA |
| JQL34 | Root | HM034 | None | 0 | NA | NA | NA | NA |
| JQL35 | Root | HM034 | None | 0 | NA | NA | NA | NA |
| JQL36 | Root | HM034 | None | 0 | NA | NA | NA | NA |

**Table S3.** Statistics associated with co-expression networks built from different tissue types.

| Network Name | Mt_General | Mt_Leaf | Mt_Nodule | Mt_Root | Mt_JQL | Mt_JQL_Nodule |
| --- | --- | --- | --- | --- | --- | --- |
| **Tissue type(s)** | Leaf, Root Nodule | Leaf | Nodule | Root | Root and Nodule | Nodule |
| **Samples** | 102 | 45 | 37 | 20 | 36 | 24 |
| **Genes included** | 24,067 | 21,822 | 21,054 | 23,773 | 23,131 | 22,123 |
| **Edges** | 289,598,211 | 238,088,931 | 221,624,931 | 282,565,878 | 267,510,015 | 244,702,503 |

